# Mechanistic Insights into Alcohol-Induced DNA Crosslink Repair by the Fanconi Anemia Nuclease Slx4-Xpf-Ercc1

**DOI:** 10.1101/2024.10.14.618148

**Authors:** Jana Havlikova, Milan Dejmek, Andrea Huskova, Anthony Allan, Evzen Boura, Radim Nencka, Jan Silhan

## Abstract

During cell division, DNA replication stalls upon alcohol-derived interstrand crosslink, employing DNA repair pathways. Acetaldehyde, a toxic metabolite of alcohol, can induce DNA crosslinks between complementary DNA strands, causing significant genomic instability. The repair of acetaldehyde DNA crosslinks (AA-ICLs) is vital for maintaining genome integrity and preventing mutagenic events. The Fanconi anemia (FA) repair pathway’s involvement in fixing the AA-ICLs is crucial, ensuring cellular homeostasis and safeguarding genomic stability. Dysregulation of the FA pathway has been linked to Fanconi anemia, a rare genetic disorder characterized by hypersensitivity to DNA-damaging agents, including acetaldehyde. However, the precise mechanism of the repair and incision of AA-ICLs is unknown. Here, we demonstrate the role of the FA pathway nuclease SLX4-XPF-ERCC1 (SXE) in the repair of AA-ICL. We have generated site-specific AA-ICL within the replication fork, and we found SXE to excise the crosslink from the replication fork, bringing evidence of its role in the repair of alcohol-induced DNA lesions. This nuclease complex performs two precise incisions around the lesion. We also demonstrate that the SXE nuclease excises an abasic site interstrand crosslink in a similar manner. Given the evidence in conjunction with previous repair studies that have been conducted, our work suggests that SXE is a versatile nuclease complex.

## Introduction

Ethanol, commonly referred to as alcohol, has been a widely consumed recreational substance for centuries. However, over the years, its use has been associated with adverse health effects. Extensive research has linked alcohol consumption to over sixty diseases, including liver damage, heart disease, cancer, neurodegenerative conditions, and mental illnesses (1). Notably, alcohol consumption has been linked to the development of several types of cancers, particularly within the digestive system. These include cancers of the oral cavity, pharynx, larynx, oesophagus, colorectal region, and liver (2,3).

Alcohol metabolism involves multiple enzymes, with the χ isoform of alcohol dehydrogenase (ADH) primarily responsible for converting alcohol to acetaldehyde. Additionally, cytochrome P450 2E1, a membrane protein expressed in the liver and catalase in microsomes, contributes to this metabolic process, even as early as in the stomach (4,5). Several studies have reported elevated levels of acetaldehyde in saliva, indicating that a portion of alcohol undergoes metabolism as soon as it enters the oral cavity (6–8). The liver contains the highest concentration of enzymes involved in alcohol metabolism, including acetaldehyde dehydrogenase (ALDH1&2), which plays a crucial role in detoxifying acetaldehyde into acetate (9). There are seven genes for alcohol dehydrogenases and 19 genes for aldehyde dehydrogenases in the human genome (10,11). However, the enzymes ALDH2 and ADH5 are considered to be the most significant for aldehyde metabolism (12,13) (Figure 1A). ALDH2-deficient alcoholics displayed increased DNA damage, outlining the ALDH2’s role in protecting genomic integrity (14).

Acetaldehyde possesses electrophilic properties that make it an exceptionally reactive compound, interacting with nucleophilic groups of amino and thiol groups alike. It forms covalent bonds with various proteins (tubulin, hemoglobin, lipoproteins, albumin, collagen), as well as with enzymes, microtubules, and DNA, leading to the disruption of cell integrity (14,15). Furthermore, acetaldehyde reacts with neurotransmitters such as dopamine to form the compound salsolinol. Salsolinol influences brain centres associated with pleasure and reward, potentially contributing to the development of alcohol addiction (15).

Importantly, acetaldehyde can react with the C-2 amino group of deoxyguanosine (dG) in DNA, forming an *N*^2^-ethylidene-2’-dG. It can subsequently form an intrastrand crosslink with the adjacent dG (16). Subsequently, the reaction with an additional acetaldehyde molecule leads to the formation of α-CH3-γ-OH-*N*^2^-propano-2’-deoxyguanosine. This lesion has a closed and open form, the latter containing an aldehyde group capable of spontaneously forming further DNA interstrand crosslinks (ICL), primarily with opposing guanine but also with proteins (17,18) (Figure 1B). Repairing DNA ICLs is essential for maintaining genomic integrity and preventing deleterious effects, such as cell cycle arrest, cell death, or cancer . The inability to effectively repair ICLs is a characteristic feature of a genetic disease called Fanconi anemia (FA) (17,19).

**Figure 1.**
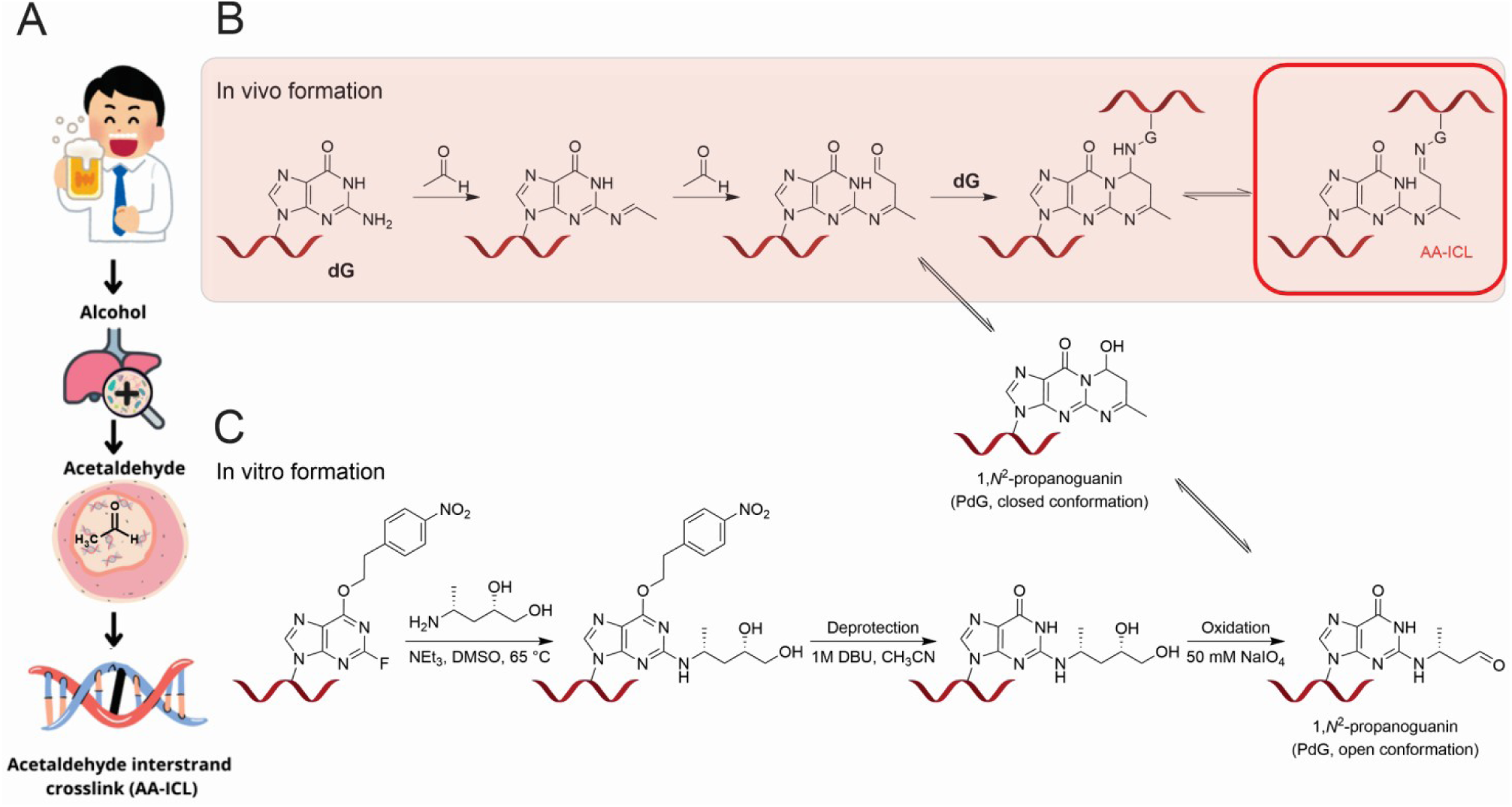
Metabolism of alcohol and formation of acetaldehyde interstrand crosslink (AA-ICL). (A) Consumption of alcohol leads to acetaldehyde formation, which can form AA-ICL in cells. (B) Formation of native AA-ICL in vivo. Two molecules of acetaldehyde react with guanosine by forming (*R*)-α-CH3-γ-OH-1,*N*^2^-propano-2’-deoxyguanosine (PdG) which thus can form a covalent bond with another guanosine on the opposite DNA strand. (C) Modification (substitution with 4-(*R*)-aminopentane-1,2-diol) of oligonucleotide immobilized on matrix to form a site-specific AA-ICL.

FA is a rare, predominantly autosomal recessive disease that results in bone marrow failure, developmental anomalies, and an increased risk of both hematologic and nonhematologic malignancies (19,20). The hypersensitivity of cells from FA patients to crosslinking agents led to the identification of the FA crosslink repair pathway, which is crucial for repairing ICLs during DNA replication. This discovery highlights the key role of the FA pathway in DNA repair and underscores its association with the maintenance of genomic integrity (21).

When the replication fork encounters a DNA crosslink during replication, it stalls, and ATR kinase is activated, employing various factors to repair the ICL (22,23). The FA pathway is initated during the unloading of the replicative helicase, leading to the activation of the FA core complex, a large E3 ligase responsible for the ubiquitylation of FANCD2 (24). This modification is a hallmark of FA and is crucial for the subsequent excision of the ICL and the initiation of further repair processes (25). The precise nuclease incisions are carried out once the DNA polymerase has progressed past the -1 position relative to the crosslink (22). For ICLs induced by nitrogen mustard, the FA nuclease complex SLX4-XPF-ERCC1 (SXE) is responsible for their excision (26). It has been shown that SLX4 stimulates the activity and specificity of the XPF-ERCC1 nuclease approximately 100 times towards replication-like structures. The research has successfully showcased that the SXE nuclease executes two precise incisions, mirroring the observations made in *Xenopus* egg extracts (26). Previously, the involvement of the FA pathway in the repair of alcohol-derived crosslinks has been elucidated, particularly in addressing acetaldehyde toxicity. Notably, FA-deficient cell lines exhibit elevated markers of DNA damage.

Moreover, the study revealed significantly reduced survival of SLX4-deficient cells (27). Further genetic experiments have strengthened the evidence and the involvement of the FA pathway in repairing acetaldehyde crosslinks. Elevated markers for DNA damage were evident in FA-deficient cell lines. The acetaldehyde exposure resulted in a significant reduction of the survival of SLX4-deficient, as well as other FA-deficient cells, derived from blood lineage progenitors (27).

The link between alcohol and cancer has been the focus of research for over a century, with recent studies beginning to unravel the molecular mechanisms involved. Seminal work has explored how alcohol, especially acetaldehyde, affects DNA damage in the context of the FA repair pathway. The study demonstrated that cells deficient in this crosslink repair pathway exhibit hypersensitivity to acetaldehyde, linking acetaldehyde exposure to DNA damage-related diseases. Furthermore, the research highlighted ethanol’s adverse effects on in utero development, hematopoiesis, and its potential teratogenicity (28).

In subsequent work, we synthesized a site-specific alcohol-induced acetaldehyde ICL (AA-ICL). Repair studies using *Xenopus* egg extracts demonstrated that the FA pathway primarily mediates the repair of these ICLs, although a secondary, alternative repair mechanism was also identified (29). More recently, a reduced form of AA-ICL was synthesized by another route, resulting in all imine bonds of the Schiff bases between the crosslink and DNA being in their reduced form. Data from this study strongly suggest that the reduced form of ICLs is exclusively repaired by the FA pathway, which was later confirmed by another study (30). However, the molecular details of how this crosslink is removed remain unclear.

Here, we focus on unravelling the intricate molecular details surrounding AA-ICL and its subsequent repair by the SXE complex. Specifically, we embarked on synthesizing a site-specific acetaldehyde crosslink (Figure 1C). Our AA-ICL specifically links opposite DNA strands within the duplex of a replication fork structure, mimicking a stalled replication scenario in the presence of the crosslink. Through meticulous enzymatic assays involving SXE, we aimed to elucidate the incision mechanism employed by these enzymes on the AA-ICL substrate. Notably, our results reveal the efficiency of SXE in cleaving AA-ICL. To broaden our investigation, we extended our analysis to the NEIL3 glycosylase, a known player in the repair of other ICLs and DNA lesions.

## Materials and Methods

### DNA oligonucleotides preparation

All DNA oligonucleotides were synthesized commercially (Eurofins Genomics). The sequences of all DNA oligonucleotides and structures of annealed substrates are provided in the supplementary file (Figure S2). Fluorescently labelled oligonucleotides were purified by band excision from 15% PAGE gel, followed by elution into TE buffer (10 mM Tris pH 8.0, 1 mM EDTA). Purified samples of ssDNA were subsequently concentrated on Amicon 0.5 with molecular cutoff 3kDa (Cytiva MicroSpinTM G-25) to approx. 10 uM.

### Synthesis of aminopentadiol

The synthesis of (4*R*)-4-aminopentane-1,2-diol has been described previously (29). Briefly: First, the commercially available (*S*)-pent-4-en-2-ol was treated with phthalimide under the Mitsunobu conditions to afford the protected (*R*)-pent-4-en-2-amine in 75-85 % yield. The phthalimide protection was exchanged for a benzyloxycarbonyl (Cbz) in a one-pot, three-step sequence affording the Cbz-protected intermediate in 56-63 % yield. The terminal double bond was dihydroxylated using the RuCl_3_-CeCl_3_ catalytic system and sodium periodate as an oxidating agent. The resulting diol (49 % yield) was isolated as a diastereomeric mixture (ca 1:1). Cleavage of the Cbz protecting group was achieved hydrogenolytically on a Pearlman catalyst. No chromatographic purification of the free amine was necessary in this final step. The overall yield of the synthesis was slightly lower than reported in the literature. Achieved purity was > 95 % (NMR-based). The scheme of the synthesis is provided in the supplementary file (Figure S2).

### Preparation of AA-ICL on fluorescently labelled substrate

The following procedure was performed according to our previous work. Briefly, a 35 NK oligonucleotide (ATGCCTGCACGAATTAAC[2-F-dI-CE]GATTCGTAATCATGGT) immobilized on solid support was incubated with (4*R*)-4-aminopentane-1,2-diol prepared as described above. The O-6 protecting group was removed by reaction with DBU. The remaining protecting groups of the oligonucleotide were removed by a 28% aqueous solution of ammonia, and the oligonucleotide was eluted from the solid support. The aqueous solution was rapidly frozen in liquid nitrogen and freeze-dried using a benchtop lyophilization system FreeZone Plus 2.5 Plus (Labconco Corporation). The sample was subsequently dissolved in 1 dM TEAAc, purified by HPLC on a semi-preparative column (Phenomenex Luna 5 μm C18 column (150 mm **×** 10 mm) and re-lyophilized. The vicinal diol was oxidatively cleaved with 50 mM sodium periodate. The aqueous solution was rapidly frozen in liquid nitrogen, freeze-dried, and purified on HPLC to afford the final oligonucleotide. The identity of the product was validated by MS (Figure S2).

The modified DNA oligonucleotide (5’-ATGCCTGCACGAATTAACG*GATTCGTAATCATGGT3’), containing the (*R*)-α-CH3-γ-OH-1,*N*^2^-propano-2’-deoxyguanosine (G*), was mixed with a partially complementary, fluorescently labelled DNA oligonucleotide (Figure 2A, sequence details in Figure S2). The mixture was annealed by slow cooling from 95°C in the substrate buffer (20 mM HEPES pH 7.0, 150 mM NaCl). The reaction mixture was incubated at 37°C for several days, followed by the isolation of DNA duplex with the ICL from a 15% denaturing PAGE gel.

### Abasic interstrand crosslink formation and isolation

Abasic interstrand crosslink (Ap-ICL) within the DNA duplex was prepared and isolated as described previously (31). Briefly, labelled and unlabelled complementary oligonucleotides were combined in equimolar ratios in a buffer (20 mM HEPES, pH 6.5, 140 mM NaCl, 0.5 mM TCEP, and 5% glycerol). The reaction mixture was annealed by heating to 95°C followed by gradual cooling to ambient temperature. To generate the Ap site, 0.5 units of uracil-DNA glycosylase (UDG) (New England Biolabs) was introduced and incubated at room temperature for 5 minutes. The resulting DNA fork containing the Ap site was incubated at 37°C to crosslink formation. The Ap-ICL was isolated from the polyacrylamide gel using a modified electrophoretic band excision protocol (32).

### Cloning, expression and purification of recombinant SLX4-XPF-ERCC1

The mouse genes for *Slx4*(1–758) and *Ercc1*(HT-3C-Full length) were cloned to pAcebac1 vectors, and the mouse gene for *Xpf* (full length) was cloned to pIDC vector. The XPF and ERCC1 constructs for expression of XPF and ERCC1 proteins were fused by Cre recombinase (New England Biolabs). The constructs for SLX4 and XPF-ERCC1 were then transformed to *E. coli* DH10EMBacY (Genova Biotech), and isolated bacmids were transfected to Sf9 cells using Fugene6 and 24-well plates with 1x10∧6 cells. Secondary recombinant baculovirus was used to *co*-infect Sf9 insect cells (2 L) at a density of 2-3x10∧6 cells/ml, and the culture was grown for 68 hours before being harvested. All purification steps were carried out in a buffer containing 20 mM Tris pH 8.0, 150 mM – 1 M NaCl, 10% (*v*/*v*) glycerol, and 3 mM B-ME. Cells were homogenized by sonication, followed by affinity chromatography using an MBP-tagged domain. Proteins were eluted with the buffer supplemented with 20 mM maltose. The complexes were then loaded onto a HiTrap™ Heparin HP 5 ml column (GE Healthcare) and eluted using a NaCl gradient. The MBP tag was cleaved overnight at 4 °C using TEV protease, while the HT tag was cleaved using 3C protease. Concentrated samples were further purified on a Superose 6 Increase 10/300 GL column (Cytiva), and the combined fractions were flash-frozen. XPF point mutants (primers for mutation are provided in Figure S1) in the XE and SXE complexes were purified using a procedure identical to the wild type.

### Cloning, expression and purification of NEIL3

Cloning, expression and purification processes were described previously (33). Briefly, the NEI domain was cloned into a modified pET-24b vector that includes the C-terminal 3C protease (HRV) site and subsequent His6x tag. The plasmid was transformed into Escherichia coli BL21 StarTM (DE3) cells (ThermoFisher). An initial 5 ml culture was grown overnight in LB medium at 37°C. Protein expression was conducted in ZY5052 autoinduction media supplemented with 50 µM ZnSO_4_. The culture was grown at 37°C until OD_600_ = 0.6-1; then, the temperature was lowered to 18 °C for overnight growth.

Harvested cells were disrupted via sonication in a lysis buffer comprising 20 mM Tris-HCl (pH 8.0), 300 mM NaCl, 30 mM imidazole pH 8.0, 10 % (*v*/*v*) glycerol and 1 mM TCEP. The lysate was fractionated by centrifugation, and the supernatant was incubated with Ni-NTA resin (Machery-Nagel) using the batch technique. Protein was eluted using an imidazole-enriched buffer.

Protein was desalted using a HiPrep 26/10 desalting column, followed by fractionation via cation exchange chromatography on a HiTrap SP HP column. The mobile phase consisted of 20 mM Tris-Hcl pH 8.0, 70 mM NaCl, 10% (*v*/*v*) glycerol and 2 mM B-ME. Protein was eluted using a NaCl gradient. Final purification was achieved via size-exclusion chromatography using a Superdex 75 Increase GL HiLoad 10/300 column equilibrated with the aforementioned buffer.

Protein purity was verified by SDS-PAGE on a 15% polyacrylamide gel. The purified NEI domain was concentrated, flash-frozen in liquid nitrogen and stored at -80°C.

### Kinetics of AA-ICL formation and stability

The reaction mixture was incubated at 37°C under near-physiological conditions (20 mM HEPES, pH 7.0; 150 mM NaCl) for several days. Periodic aliquots were taken for stability assays. Afterward, DNA duplexes containing ICLs were purified by isolatation from 15% denaturing PAGE gel. The isolated ICLs were diluted in substrate buffer and subjected to a stability assay at 37°C the aliquots were taken in given timepoints. Subsequently, the aliquots were analyzed using a 15% denaturing gel.

### Nuclease assay

All reactions were carried out in a nuclease buffer (25 mM Tris pH 8.0, 50 mM NaCl, 2 mM MgCl_2_, 1 mM TCEP, 0.1 mg/ml BSA, 5% glycerol) at 25°C. The crosslinked substrate was synthesized as described above. Reactions were initiated by mixing 40 nM of the given substrate with 100 nM of the enzyme SXE. After the incubation period, the reactions were quenched with 80% formamide, 200 mM NaCl, 10 mM EDTA, 0.01 % bromophenol blue and analyzed on 15% denaturing PAGE gel. The gels were visualized using the Amersham Typhoon laser scanner (Cytiva). The signals were analyzed using ImageQuant TL version 8.2.0 software (Cytiva).

## Results

### Preparation of site-specific alcohol-induced interstrand crosslink

The synthetic crosslink in this study is chemically identical to the alcohol-induced interstrand crosslink (AA-ICL) that forms in DNA following alcohol consumption. A synthetic single-stranded DNA oligonucleotide, modified at a specific site with the 2-InoF residue, was coupled with (4*R*)-4-aminopentan-1,2-diol via an SNAr reaction. In this reaction, the C-2 fluorine atom of the 2-InoF residue is easily replaced by a nucleophile, such as a primary amine. The resulting product serves as the starting material for producing a chemically identical lesion to that formed by the reaction of two acetaldehyde molecules with guanine within DNA. The 1,2-diol was converted to PdG through oxidation with NaIO_4_. The resulting product was then purified using HPLC. The appropriate fractions were concentrated and used in subsequent crosslinking reactions (Figure 2A).

Four different substrates generated from the original sequence were prepared to fully address the enzymatic cleavage of AA-ICL by SXE. These substrates contained an identical single-stranded DNA strand with the PdG residue prepared for crosslinking. The opposing strand was designed so the annealed oligonucleotide resembled a DNA replication fork in the shape of the letter Y. In each case, a fluorescent dye was placed at one end of the replication fork to monitor the reaction. Therefore, two 3′ and two 5′ labelled oligonucleotides were synthesized (Figure 2A).

### Formation and stability of AA-ICL, the rate of formation of AA-ICL is relatively slow, but so is its degradation

The formation of DNA crosslinks was observed in different oligonucleotides containing the naturally identical lesion PdG (Figure 2A). The oligonucleotides were annealed, and the crosslinking reaction was carried out in the dark. Subsamples were taken at selected intervals to determine the percentage of AA-ICL formed within the reaction mixture. The reaction was resolved on a 15% denaturing PAGE gel, and the reaction rate was 0.1% per day (Figure 2B,C). To revalidate that the crosslink forms specifically between PdG and the opposing dG, the dG was replaced with inosine (29). The crosslink was formed only within the duplex containing PdG with the opposing dG. In the case of the inosine-containing oligonucleotide, no crosslink was observed, confirming the necessity of the C-2 amino group (Supplementary Figure S2).

After the crosslinking reaction, the AA-ICL was purified from the gel. To address the stability of AA-ICL, the purified crosslinked DNA was allowed to degrade over time at 37°C. The reactions were analyzed on a denaturing gel, similar to the method used for formation analysis. From this experiment, it was evident that degradation is again a relatively slow process, and the reaction does not reach full conversion. It appears that the crosslink forms an equilibrium with uncrosslinked DNA.

**Figure 2.**
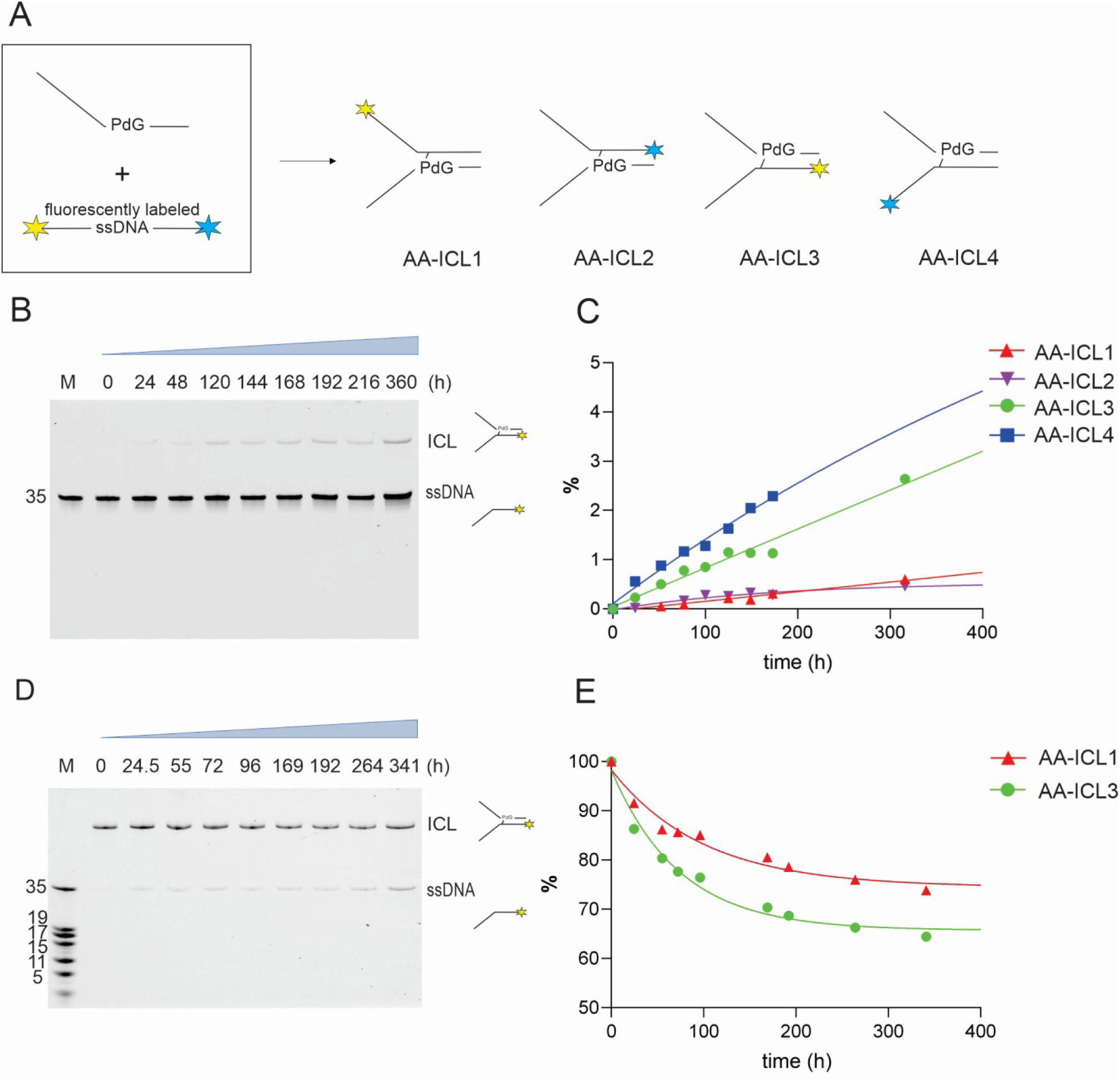
AA-ICL formation and stability rate. (A) Representation of DNA replication fork substrates used in this study differing in the position of the ICL site and its polarity. (B) 15% denaturing PAGE gel showing AA-ICL formation from annealed DNA fork. (C) The gel bands corresponding to the reactants (ssDNA) and the products of AA-ICL formation (ICL) were quantified. The relative ratios of ICL formation were plotted against reaction time for various uncrosslinked AA-ICL substrates. (D) Degradation of AA-ICL into ssDNA was monitored over time at 37°C using a 15% denaturing PAGE gel. (E) The relative proportion of degraded AA-ICL was plotted against reaction time for various AA-ICL substrates (normalized to 100%).

### Fanconi anemia enzyme SXE cleaves alcohol-derived acetaldehyde DNA interstrand crosslink

To investigate the role and molecular mechanism of the FA enzyme complex SXE in DNA repair, uncrosslinked and crosslinked DNA forks containing site-specific AA-ICLs were subjected to enzymatic reactions. The DNA fork AA-ICL substrates contained the fluorescent label on the top strand of the left-handed fork either on the 5′ or 3′ end, and the reactions were resolved on 15% denaturing PAGE gel (Figure 3). In qualitative reaction, cleavage of the DNA resulted in two distinct bands that migrated at higher molecular mass than the 35 nt arm of the uncrosslinked fork (Figure 3B). This indicated that a DNA crosslink (AA-ICL1) was excised on the top strand as the uncrosslinked control remained unchanged by the reaction. The first incision indicated as P1 was the higher migrating band, where P2 was likely cleaved from the other side of the crosslink (Figure 3B, F). This strand was also observed in the time course of this particular reaction, where the original crosslinked AA-ICL substrate was completely digested by the enzyme within 60 min (Figure 3C).

To confirm the results, a further experiment was carried out under identical conditions with identical substrates labelled on the opposite side of the top strand, the 3′ end, of the same DNA strand (AA-ICL2). For these substrates, the stability of the probe was sensitive to degradation, and the signal decreased significantly during the substrate preparation and purification.

Consistent with the experiment above, where the fluorescent probe was placed on the 5′ end, the assay on this fork resulted in two incisions. The bands corresponding to incisions P1 and P2 appeared and migrated as larger products than the ssDNA arm of a single fork. Additionally, the enzyme nicked the fluorophore-containing 3′ end, resulting in a short non-specific PN product (Figure 3D). As before, the uncrosslinked DNA fork, where only the top strand was labelled, did not exhibit either P1 or P2 incisions except for the production of the small PN product.

Similarly to the qualitative experiments, the gels from the time-course reaction for AA-ICL exhibited both bands corresponding to incisions P1 and P2, akin to those observed for the 5’ labelled DNA fork. However, as the nicking of the 3′ end probe continued, all bands disappeared.

The nicking activity of SXE on the 3′ end, where the fluorescent probe is removed (product P3), can be explained by the P1 or P2 incisions. When the 3′ arm is cut at the dsDNA junction, it resembles a scenario where the fork has been rotated by 180°. Thus, we assume that in this case, SXE does not discriminate between the fork junction and the dsDNA containing the 3′ fluorescent probe (Figure 3E).

Our findings were confirmed when we quantified the enzyme kinetics from these measurements and plotted the rate of the repair reaction (Figure 3F). The reaction rate was determined for substrate removal using the catalytic step. The rate for the fork labelled at the 5′ end (AA-ICL1) was k = 0.1030 s^-1,^ whilst the rate for the 3′ end labelled fork (AA-ICL2) was k = 0.2527 s^-1^suggesting that the fluorescent probe on the free arm of the replicate fork might actually may affect and decrease the activity SXE.

To confirm the location of the SXE incision in the DNA fork, we placed a fluorescent dye on the bottom strand of the left-handed fork. The resulting cleavage confirmed that digestion of P1 and P2 occurred on the fork’s bottom strand, demonstrated by the formation of products smaller than the ssDNA arm. Consistently, these incisions were observed in the uncrosslinked DNA fork (Figure S3). Further revalidation revealed that these incisions were site-specific and dependent on the presence of the active SXE complex, as reactions with catalytic point mutants did not produce any of these incisions (Figure S1). This demonstrates the crucial role of SXE in recognizing and processing AA-ICL within DNA forks, providing valuable insights into its function in DNA repair mechanisms.

**Figure 3.**
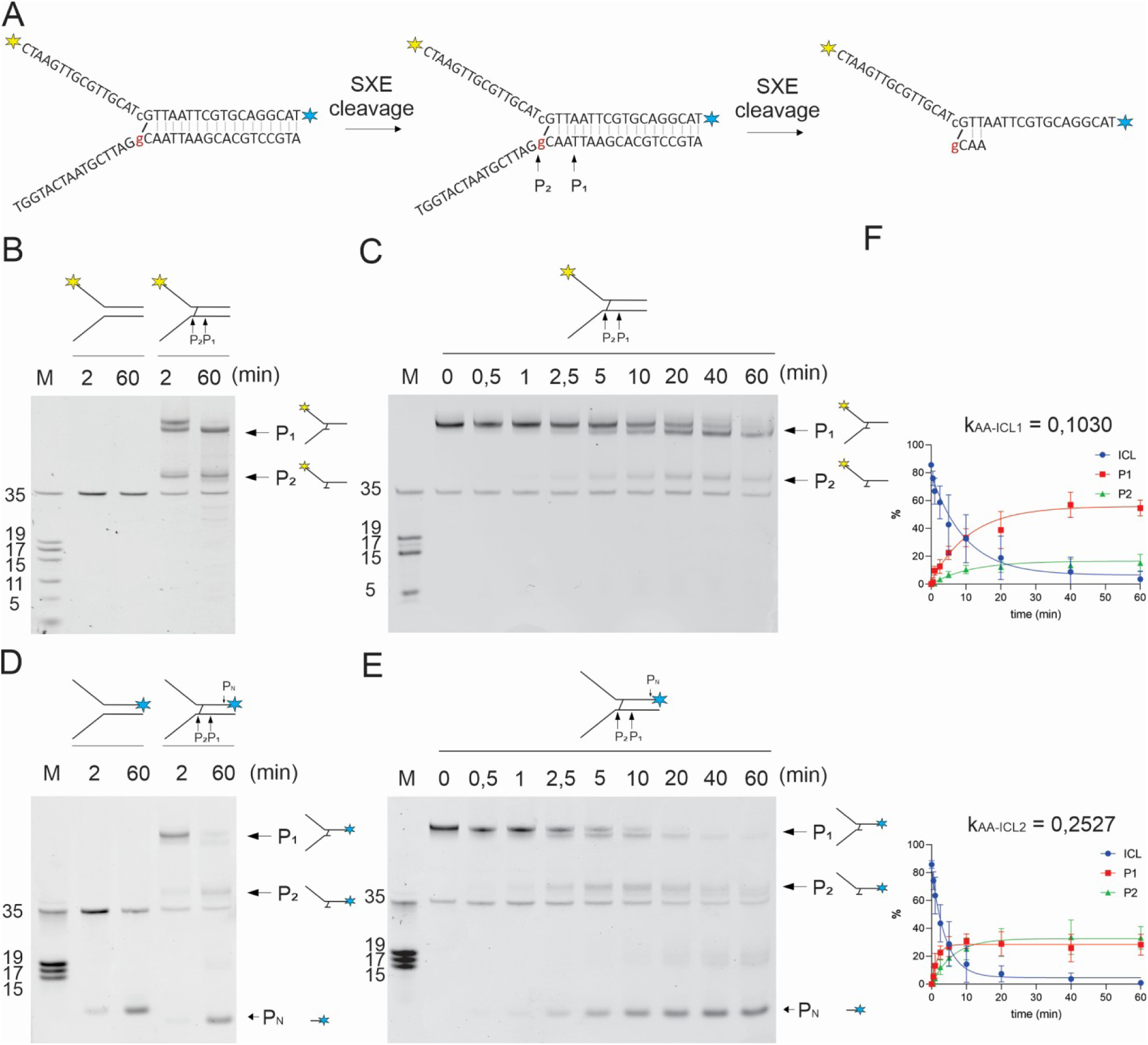
Cleavage mechanism of AA-ICL by SXE nuclease. (A) Scheme of the cleavage sites around the AA-ICL on the bottom strand of a replication fork. The substrate is fluorescently labeled either at the 5′ end (AA-ICL1) or the 3′ end (AA-ICL2) of the upper strand (not both). (B, D) Qualitative SXE nuclease assay showing non-crosslinked and crosslinked substrates, with SXE products (P_1_ and P_2_) migrating above the ssDNA control, and non-specific nicking of the fluorescent probe P_N_. (C, E) Time course of the SXE nuclease assay demonstrating the enzymatic cleavage of AA-ICL substrates fluorescently labeled at the (C) 5′ and (E) 3′ ends. (F) Data from panels C and E were plotted in a graph and fitted with exponential decay to determine reaction rates. Reactions were repeated in triplicate, and the error is represented as standard deviation (SD).

### SXE excises the abasic site interstrand crosslinks (Ap-ICLs), a substrate of NEIL3 glycosylase

Given that SXE has been shown to cleave both AA-ICLs (this study) and nitrogen mustard ICLs (26), we investigated how SXE handles abasic site interstrand crosslinks (Ap-ICLs). Ap-ICLs are formed through aldehyde crosslinking with the opposite base, originating spontaneously from abasic sites (34) (Figure 3A). These crosslinks are analogous to AA-ICLs (Figure 4). It is important to note that the structural orientation and directionality of Ap-ICLs within the DNA duplex are opposite to those of AA-ICLs.

In SXE nuclease assays, we qualitatively compared the incisions of Ap-ICL1 with AA-ICL3 on left-handed forks, with the fluorescent probe located on the bottom strand (Figure 4B). The control reaction with AA-ICL produced the two expected products (P1 and P2), consistent with previous results (Figure S3). Similarly, Ap-ICL DNA substrates also yielded products P1 and P2, where P1 corresponds to the incision of the first part of the crosslink and P2 corresponds to the removal of the fluorescently labelled DNA strand, either on the 3′ or 5′ end of the crosslink, completing the excision step.

These experiments demonstrate that Ap-ICL is a substrate for SXE nuclease. Despite the differences in crosslink orientation compared to AA-ICL, SXE effectively cleaves Ap-ICLs similarly. This indicates that SXE can process a range of DNA crosslinks, including those formed from spontaneously occurring abasic sites.

### Mouse NEIL3 enzyme responsible for Ap-ICL repair does not cleave AA-ICL

Since the Ap-ICL is repaired by NEIL3 glycosylase in an upstream event before employment of the FA pathway, we were interested in how NEIL3 processes AA-ICL. We have employed the enzymatic domain NEI responsible for removing Ap-ICL in replication-coupled repair. Under the same reaction conditions previously used for testing SXE, we compared Ap-ICLs with AA-ICLs. The substrates (Ap-ICL2, AA-ICL1) were selected to favor NEIL3 glycosylase activity, as the NEI domain requires single-stranded DNA (ssDNA) at the 3′ end.

Unlike SXE, the NEI domain of NEIL3 was incapable of excising or reverting the AA-ICL DNA fork substrate. On the other hand, the Ap-ICL-containing fork was expectedly cleaved by glycosylase activity and served as a control to validate the activity of our NEI enzyme in this assay’s conditions.

Overall, we have demonstrated that SXE can cleave crosslinks that may arise as a result of alcohol consumption, which elevates the levels of acetaldehyde and are relatively stable. SXE excises this acetaldehyde crosslink mainly with two distinct incisions within the DNA strand of replication fork mimetics while leaving the opposite strand intact. Moreover, Ap-ICL is also recognized and cleaved by SXE. This data demonstrates that SXE cleaves not only AA-ICL but also Ap-ICL and other crosslinks, such as nitrogen mustard, as we demonstrated previously (26).

**Figure 4.**
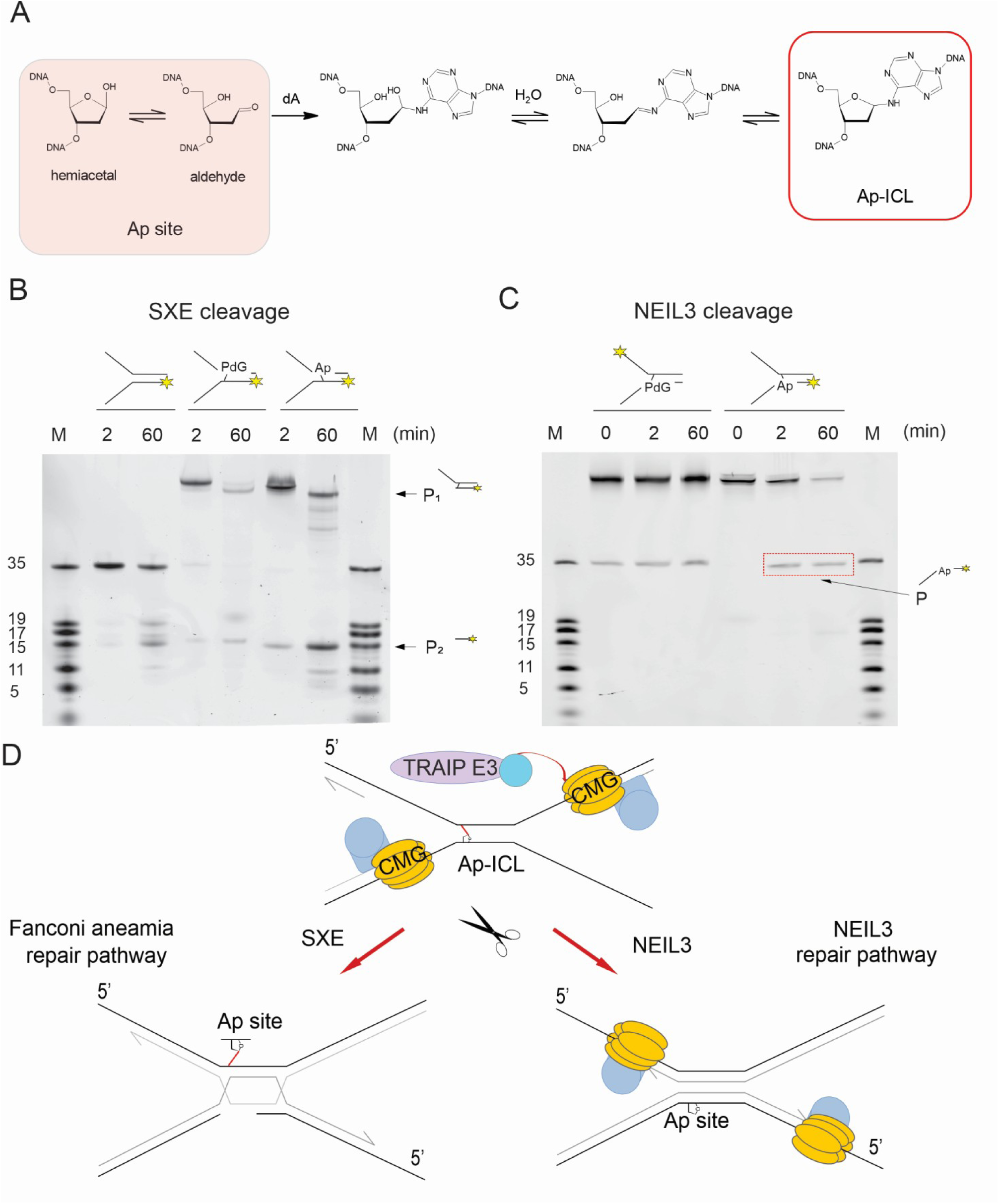
Comparison of cleavage mechanism of AA-ICL and Ap-ICL. (A) The model of the formation of Ap-ICL in vivo. (B) SXE nuclease assay of AA-ICL (AA-ICL3) compared to Ap-ICL (Ap-ICL1) with respective nuclease products P_1_ and P_2_. (C) NEIL3 glycosylase assay of the AA-ICL (AA-ICL1) and Ap-ICL Ap-ICL2 by catalytic NEI domain of mouse NEIL3 glycosylase. NEIL3 enzymatic product is ssDNA product visible only for Ap-ICL substrate. (D) A schematic model of Ap-ICL cleavage and potential repair mechanisms. Both NEIL3 and the FA repair pathway recognize and process Ap-ICL. In the FA pathway, Ap-ICL is cleaved similarly to AA-ICL by the SXE nuclease, resulting in double-strand breaks that serve as substrates for further repair steps, such as homologous recombination. In contrast, during the NEIL3 repair pathway, the helicase is not unloaded from the DNA, and NEIL3 unhooks the Ap-ICL.

### Summary

Our results demonstrate that SXE can cleave acetaldehyde interstrand crosslinks (AA-ICLs) resulting from alcohol consumption. SXE excises AA-ICL primarily through two distinct incisions within the DNA strand of replication fork mimetics while leaving the opposite strand intact. Additionally, SXE recognizes and cleaves Ap-ICL. Thus, SXE is a versatile nuclease that targets the DNA structure rather than being selective for a specific type of crosslink. This underscores the FA pathway’s role as a universal repair mechanism for a wide range of challenging crosslinks.

## Discussion

### The formation of the AA-ICL

The formation of AA-ICLs presents a significant challenge in understanding alcohol-related DNA damage. This is especially pronounced in tissues frequently exposed to alcohol, such as the liver and oesophagus, where chronic exposure can overwhelm DNA repair mechanisms and contribute to cancer development.

In addition to cellular metabolism, acetaldehyde is produced directly in the oral cavity through the microbial breakdown of alcohol (6). Studies with germ-free mice have shown lower acetaldehyde levels in the digestive tract compared to microbiota-colonized mice, suggesting that microbial fermentation may significantly influence gastrointestinal cancer risk in alcohol consumers (35). While acetaldehyde is predominantly known as the primary oxidative metabolite of ethanol within the human organism, it also arises naturally under physiological circumstances through the metabolism of threonine, β-alanine, deoxyribose phosphate, or sugar metabolism (36). The pervasive generation of acetaldehyde through its consumption, microbial, and metabolic processes underscores the importance of understanding and improving DNA repair mechanisms for acetaldehyde-induced damage, like AA-ICLs, to reduce cancer risk in alcohol-exposed tissues.

### The possible implications of alcohol-induced DNA damage and its impact on genomic stability

The exact number of AA-ICLs formed during alcohol consumption can vary depending on several factors, including the amount of alcohol consumed, the efficiency of acetaldehyde detoxification pathways, such as those involving aldehyde dehydrogenase, and individual genetic differences. However, quantifying the precise number of AA-ICLs formed in a typical scenario is challenging and typically cannot be provided in straightforward figures. The extent of AA-ICL formation can be inferred from studies examining the levels of acetaldehyde-DNA adducts or the crosslinking frequency in cells exposed to acetaldehyde. However, these studies typically express the results regarding adducts per million nucleotides or the frequency of crosslinking events rather than an absolute number of crosslinks formed.

Here, we present the percentage of AA-ICL formation at a site-specific position within a DNA replication fork structure formed in vitro. The data reflect the extent of AA-ICL formation following the synthesis of the primary precursor, PdG. Consequently, the results indicate the percentage of crosslinking that occurs between an acetaldehyde-induced adduct and deoxyguanosine residue on the complementary DNA strand, particularly within a CpG sequence.

While various types of adducts resulting from acetaldehyde exposure have been identified, the PdG adduct is considered particularly mutagenic (37,38). The formation of the PdG precursor in vitro itself is relatively low; it constitutes less than 10% of the primary acetaldehyde adduct, *N*^2^-ethylidene-dG, as observed by Wang et al. (39). Under conditions of excessive acetaldehyde exposure, significantly higher than levels observed after alcohol consumption, the formation rate of PdG is approximately 1 PdG per 1.3 × 10⁸ dG per 24 hours. In the human genome, which contains approximately 1.23 × 10 ^9^ GC pairs, this would correspond to about ten molecules of PdG formed every 24 hours. Notably, the synthesis of PdG requires more than one molecule of acetaldehyde per molecule of deoxyguanosine, highlighting the inefficiency of this process in vitro (39).

However, Theruvathu et al. (2005) demonstrated that in vivo, the formation of the PdG adduct from acetaldehyde is significantly facilitated by the presence of basic molecules such as histones and polyamines, including spermidine (40). This suggests that in vivo formation is much more efficient than in vitro. These basic molecules possess amino groups and are crucial for converting acetaldehyde into reactive intermediates that lead to PdG adduct formation. While acetaldehyde alone can interact with DNA to form adducts, these basic molecules greatly enhance PdG formation efficiency (40). These reactions can result not only in the formation of AA-ICLs between DNA strands but also in the formation of DNA-protein crosslinks (18).

Due to the differences in the rates of PdG formation under in vivo and in vitro conditions and the need for an AA-ICL at a specific position, the PdG was synthesized directly at the required site in this study. Therefore, AA-ICL formation reflects its occurrence at a defined position within the replication fork. Subsequently, incubation with a partially complementary strand resulted in the formation of a DNA replication fork substrate with AA-ICL.

Despite significant differences we observed between various substrates, the formation of native AA-ICLs reached a maximum of approximately 5% after 400 hours, likely due to the system reaching an equilibrium between ICL and the DNA duplex. In comparison, Kozekov et al. (2003) reported that AA-ICL formation reached up to 38% over 21 days (41). However, their study examined AA-ICL formation in 12-mer and 15-mer duplexes, whereas our research investigated AA-ICL formation in a Y-shaped substrate. Also, our analysis of multiple substrates suggests that local sequence context may play a role in the rate of AA-ICL formation, potentially due to differences in GC content. This is similar to what has been observed with Ap-ICL formation (31).

Acetaldehyde typically binds to intracellular macromolecules by forming imines, or Schiff bases, which are often unstable and can readily revert to their original compounds (42). In this study, we demonstrated both the formation and decomposition rates of the AA-ICL. The isolated AA-ICL decomposed and reached equilibrium at approximately 70% of the original substrate after 400 hours, indicating that the AA-ICL is relatively stable. The partial reversal of the crosslinking process is potentially influenced by the interconversion between the open and closed forms of PdG (41).

### AA-ICL incision and how does it compare to what was observed

As we have previously demonstrated on nitrogen mustard ICL, the SXE nuclease is capable of cleaving ICLs through a double excision mechanism, making two precise cuts on the one strand of the stalled replication fork (26,43). This cleavage pattern was observed on both non-crosslinked controls and ICLs, indicating cleavage at two sites on the complementary strand. These findings support the hypothesis that SXE-mediated cleavage results in the unhooking of the ICL.

Although Hodskinson et al. (2020) demonstrated that AA-ICLs are repaired in vivo by SXE and another unknown mechanism, the precise molecular mechanism remains unclear (29). Here, we propose such a mechanism, which involves two sequential cuts around the AA-ICL only in the bottom strand with a 3′-ssDNA arm of the replication fork: the first at the 3′ end and the second at the 5′ end of the minor strand. Notably, the similar rates at which these cuts occur suggest that SXE acts indiscriminately on our DNA fork substrates. However, under in vivo conditions, factors such as other FA proteins or the X-structure itself may influence the preference and specificity of the AA-ICL incision. These incisions unhook the AA-ICL and separate the parental DNA duplex strands, allowing the damaged strands to undergo repair via the FA repair pathway. This process is vital for restoring replication and maintaining genomic stability and cellular function.

We compared our findings with studies on other types of ICLs, specifically Yperit (mustard gas and its derivatives such as nitrogen and sulfur mustards) and psoralen ICLs. In Hodskinson et al. (2014) we examined mustard gas ICLs, where SXE endonuclease made two incisions around the ICL on the lower DNA strand: first at the 3′ end and then at the 5′ end of the minor strand, which corresponds to the free end of the replication fork (26). This sequence of incisions aligns with our findings for AA-ICLs, suggesting a similar repair mechanism.

Additionally, Kuraoka et al. (2000) investigated psoralen-ICLs, which are formed under UV radiation and are repaired in different manner. They found out, that the XE nuclease cleaves the strand with a free 3′ end, initially at the 5′ end and then at the 3′ end of the lower DNA strand, with a delay between the cuts (43). The SXE complex appears to follow a similar cleavage mechanism as in psoralen-ICLs, reinforcing that SXE makes two sequential cuts on the minor strand without cleaving the main DNA strand. These insights are crucial for understanding DNA repair mechanisms for ICL-induced damage and how cells maintain genomic stability in response to various genotoxins.

We demonstrated not only that AA-ICLs are excised by SXE, but also that SXE cleaves Ap-ICLs in a similar manner, functioning as a versatile and multifunctional ICL nuclease. These findings suggest that SXE can cleave various types of ICLs, including those caused by acetaldehyde, abasic sites, sulfur mustard, and psoralen, making it a versatile tool for ICL unhooking.

### Unlike Ap-ICL, NEIL3 does not repair AA-ICL and likely acts upstream of the FA pathway

Lastly, our previous work has shown that AA-ICL repair occurs via two distinct mechanisms: the FA pathway and an as-yet-unknown mechanism (29). Here, we provide complementary data to the work in *Xenopus* egg extracts and also demonstrate that, unlike Ap-ICLs, NEIL3 does not cleave AA-ICLs. Thus, the unknown mechanism of AA-ICL repair does not involve the NEIL3 pathway. This aligns with the enzymatic properties of the specialized DNA glycosylases from the Fpg/Nei family, which form covalent adducts with damaged sites, similar to other Fpg/Nei glycosylases. SXE, on the other hand, is a nuclease that does not appear to be specific to crosslinks, as it cleaves relatively far from the damage site, regardless of the nature of the ICL. This reflects the strengths of SXE and the versatility of the FA pathway. Conversely, NEIL3 does not seem to be a universal ICL repair mechanism but rather a specialised ICL repairpathway. As described by Semlow et al. (2016) and further elaborated by Huskova et al. (2020), NEIL3 is the primary mechanism initially employed for repairing Ap-ICLs and similar lesions (33,44). When this mechanism fails, the helicase is unloaded, triggering the FA repair pathway to unhook the ICL (45–47). Therefore, the FA repair pathway is considered a universal mechanism capable of repairing all types of ICLs.

## Supplementary Information

**Figure S1.**
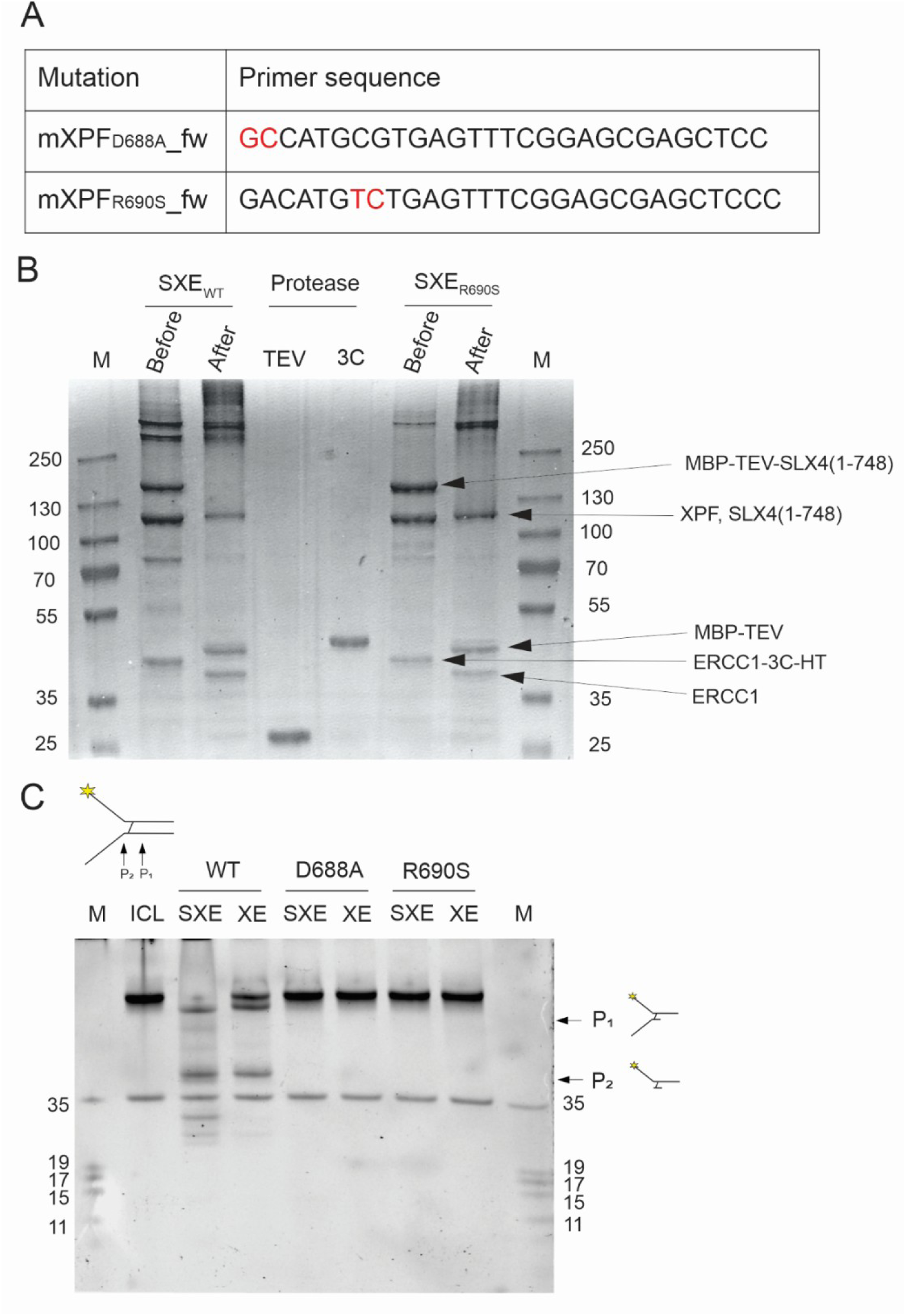
Expression and purification of SXE complex and its mutant forms. (A) Primers used for mutation. (B) Cleavage of fusion tags and purification of final protein complex. (C) Enzymatic assay of mutant forms of SXE and XE. Cleavage of AA-ICL after 60 minutes.

**Figure S2.**
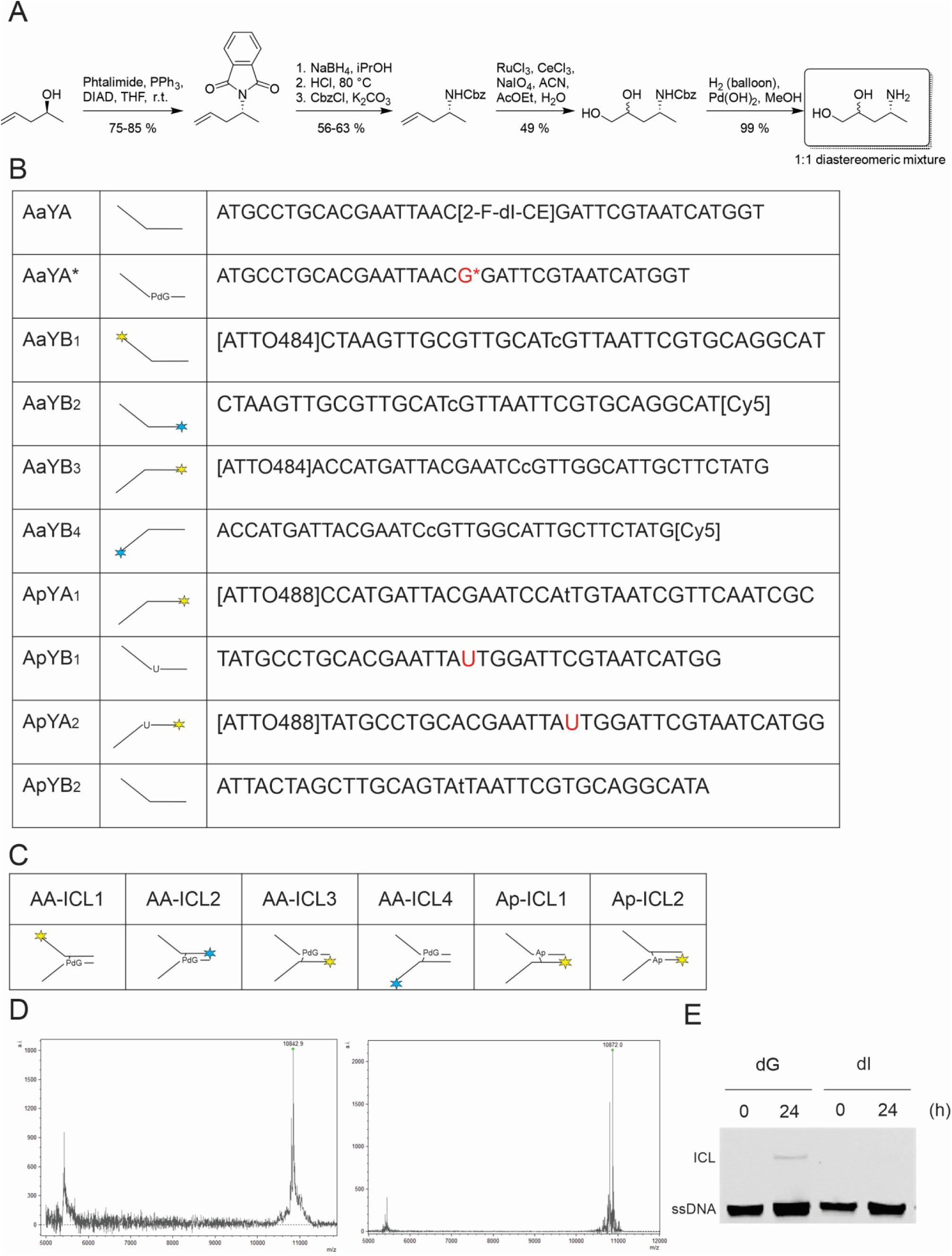
Preparation of ICL. (A) Scheme of synthesis of (4*R*)-4-aminopentane-1,2-diol. (B) Sequences of DNA oligonucleotides used in this study, where G* corresponds to (*R*)-α-CH3-γ-OH-1,*N*^2^-propano-2’-deoxyguanosine (PdG). The star’s colour refers to the fluorescent dye used (yellow - ATTO488, blue - Cyanin5). (C) Representation of prepared ICL used in this study. (D) MALDI-MS spectrum of the modified oligonucleotide before (10873,7) and after oxidation (10845,1). (E) Denaturing PAGE gel shows the formation of AA-ICL in the presence of deoxyguanosine (dG), however, in the presence of deoxyinosine (dI), no AA-ICL is formed.

**Figure S3.**
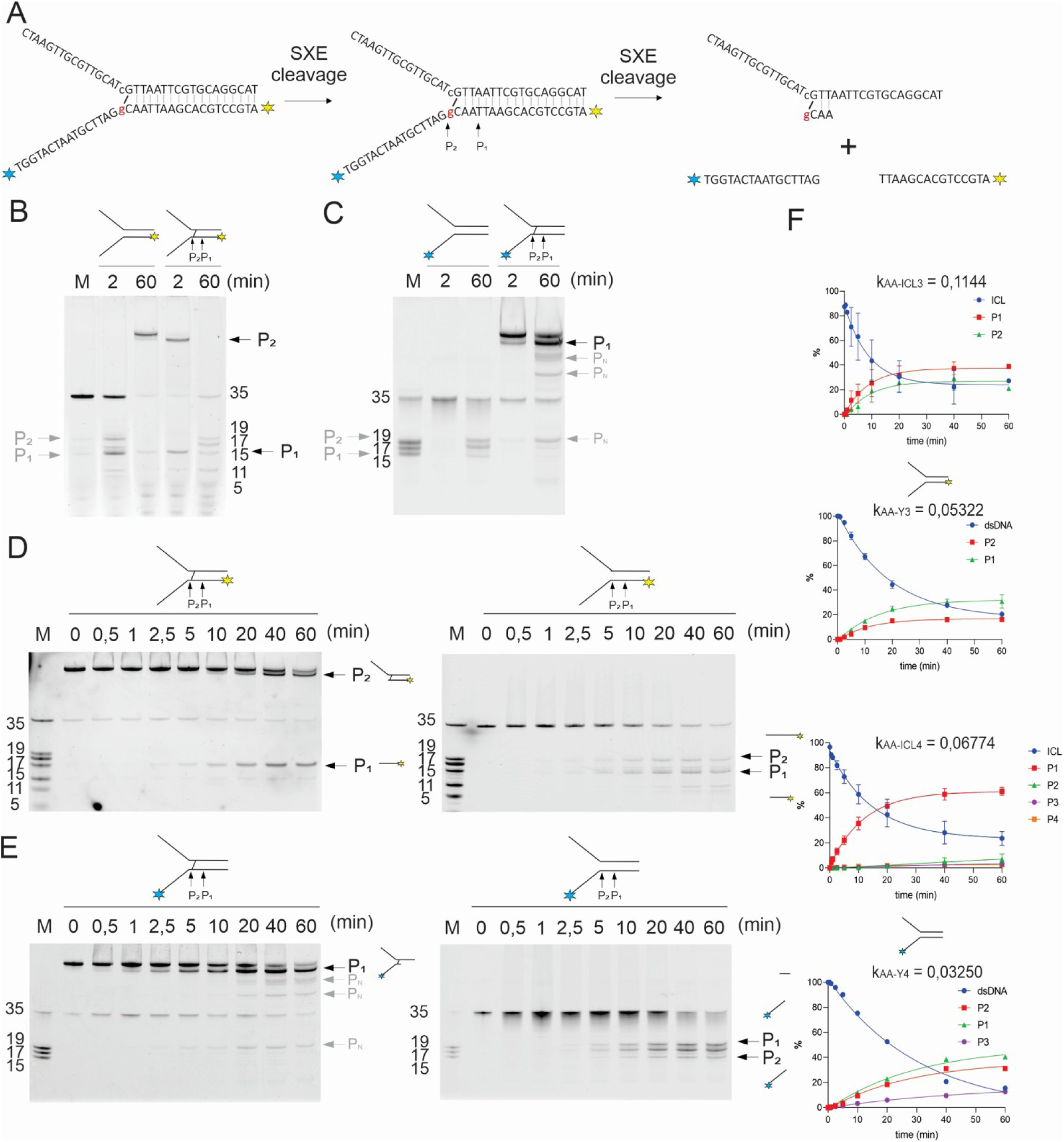
Cleavage mechanism of AA-ICL (fluorescently labelled on the bottom strand) by SXE nuclease. (A) Scheme of the cleavage sites around the AA-ICL on the bottom strand of a replication fork structure. The substrate is fluorescently labelled either on the 5′ end (AA-ICL3) or 3′ end (AA-ICL4) of the bottom strand (not both). (B, C) Non-crosslinked substrate showing cleavage on the bottom strand, resulting in two main products demonstrating the cleavage mechanism. (D, E) Kinetics of cleavage of AA-ICL substrate fluorescently labelled at 5′ end and 3′ end. (F) Graphics of AA-ICL cleavage.

